# The genome of the cryopelagic Antarctic bald notothen, *Trematomus borchgrevinki*

**DOI:** 10.1101/2024.06.17.599359

**Authors:** Niraj Rayamajhi, Angel G. Rivera-Colón, Bushra Fazal Minhas, C.-H. Christina Cheng, Julian M. Catchen

**Author notes:** Corresponding Author: Julian Catchen.

## Abstract

The Antarctic bald notothen, *Trematomus borchgrevinki* (Notothenioidae) occupies a high latitude, ice-laden environment and represents an extreme example of cold-specialization among fishes. We present the first, high quality, long-read genome of a female *T. borchgrevinki* individual comprised of 23 putative chromosomes, the largest of which is 65 megabasepairs (Mbp) in length. The total length of the genome 935.13 Mbp, composed of 2,095 scaffolds, with a scaffold N50 of 42.80 Mbp. Annotation yielded 22,567 protein coding genes while 54.75% of the genome was occupied by repetitive elements; an analysis of repeats demonstrated that an expansion occurred in recent time. Conserved synteny analysis revealed that the genome architecture of *T. borchgrevinki* is largely maintained with other members of the notothenioid clade, although several significant translocations and inversions are present, including the fusion of orthologous chromosomes 8 and 11 into a single element. This genome will serve as a cold-specialized model for comparisons to other members of the notothenioid adaptive radiation.

## Introduction

As the intensity of climate change has increased, significant scientific resources are being invested in understanding and modeling its biological effects. This prospectively-focused work seeks to understand contemporary changes in core ecological processes (Scheffers *et al*. 2016), as well as to model future effects, such as changes in species diversity and range (García Molinos *et al*. 2016). An alternative way to understand the effects of climate change is to look retrospectively, to understand how past geologic changes affected climate and how species evolved in response. One prominent area of focus is the adaptive radiation of the Antarctic notothenioid fishes.

Contemporary Antarctica is the coldest and driest continent, encircled by the Southern Ocean which maintains year-round freezing temperatures (-1.9°C) in the high-latitude continental shelves (Cziko *et al*. 2014). However, 110-90 million years ago (MYA), Antarctica was a connected land mass that exhibited a temperate climate (Zachos et al. 2001; Eastman 1993); during the late Eocene (40 MYA) the climate was cool-temperate, similar to present-day southern New Zealand and South America, and the waters surrounding the continent supported a diverse and cosmopolitan fish fauna (Eastman 2005).

Over time, tectonic forces isolated Antarctica (Storey and Granot 2021) leading to the formation of the Antarctic Circumpolar Current (ACC) which established a thermal barrier for water masses on the current’s northern and southern sides (Kennett 1977; Barker and Thomas 2004; Livermore *et al*. 2005). As Antarctica cooled, ice sheets formed and destroyed benthic habitat which, along with decreasing water temperatures, led to the extinction of nearly all ancestral Eocene fish fauna (Eastman 2005).

Despite this loss of habitat and life, a lineage of ancestral notothenioid fish evolved *in situ*, filling many newly opened niches assisted by the evolution of Anti-Freeze Glycoproteins (AFGPs) – one of two independent appearances of these types of proteins across fish taxa (Devries 1971; Chen *et al*. 1997; Zhuang *et al*. 2019). These proteins prevent the growth of ice crystals within fish enabling their survival in the glaciated Southern Ocean (DeVries and Cheng 2005). Over a relatively short span of evolutionary time (i.e., 10.7 million years), this ancestral lineage of notothenioids experienced an adaptive radiation (Eastman 2005) as they exploited the newly available ecological niches within the Southern Ocean. For example, despite the absence of swim bladders, evolving notothenioids colonized multiple water column habitats by reducing bone ossification and accumulating lipid deposits in muscle, with some species achieving neutral buoyancy (Eastman 2024).

The Antarctic clade of notothenioids, known commonly as cryonotothenioids, encompass 128 species of which 110 are endemic to the Southern Ocean where they dominate the fish fauna in species diversity and biomass (>90%) (Eastman 2005; Eastman and Eakin 2021).

Among cryonotothenioids, the bald notothen *Trematomus borchgrevinki* (Notothenioidae) represents one of the most extreme examples of cold-specialization, being the sole cryopelagic species inhabiting the coldest and iciest high-latitude waters – the ice platelet layer under surface fast ice in McMurdo Sound, Antarctica (Hunt *et al*. 2003). This species exhibits the lowest recorded blood freezing point and highest serum thermal hysteresis among studied cryonotothenioids (DeVries and Cheng 2005) and cannot tolerate elevated temperatures – *T. borchgrevinki* will succumb at approximately 6°C, the lowest incipient upper lethal temperature known for fish (Somero and DeVries 1967). Among other phenotypic changes, cold-specialized notothenioids have lost the near-universal inducible heat-shock response (Hofmann et al. 2000) and *T. borchgrevinki* is nearly silent in its transcriptional responses to heat stress (Bilyk and Cheng 2014; Bilyk *et al*. 2018) making it particularly vulnerable to a warming environment.

We present the first high quality, chromosome-level genome assembly of *T. borchgrevinki* from high-latitude McMurdo Sound. Serving as a model of cold-adapted and cold-specialized physiology, the genome resource of *T. borchgrevinki* will enable comparisons with other members of the adaptive radiation, against those that span a cline across the southern latitudes, such as *Cottoperca gobio* to *Trematomus bernacchii* (Bista *et al*. 2023), or against the large, migratory deep water predator *Dissostichus mawsoni* (Hanchet *et al*. 2008; Jae Lee *et al*. 2021), or against their highly derived, hemoglobin-less relatives, the icefishes (Kim *et al*. 2019; Rivera-Colón *et al*. 2023).

## Methods & Materials

### Species sampling and processing

A female *T. borchgrevinki* individual was sampled from McMurdo Sound, Antarctica (77.5°S, 165°E) as originally described in Rayamajhi, et al. (2022). The fish was caught by hook and line through holes drilled through annual sea ice and transported back to the aquarium facility at McMurdo Station. The fish was anesthetized using MS222 (Western Chemicals/Syndel, WA), and tissues were dissected on ice, flash frozen in liquid nitrogen and stored in a -80°C freezer until use. Fish handling and sampling complied with the University of Illinois, Urbana-Champaign (UIUC) IACUC approved protocol 17148.

### HMW DNA and long-read genome sequencing

High molecular weight (HMW) DNA was prepared from frozen muscle using the Nanobind HMW Tissue DNA Kit (Circulomics/Pacific Biosciences) following vendor instructions. The concentrations of recovered HMW DNA were determined using Qubit dsDNA Broad Range Assays on a Qubit v.3 fluorometer (Invitrogen).

PacBio continuous long read (CLR) library preparation and sequencing were carried out at the University of Oregon Genomics & Cell Characterization Core Facility. The HMW DNA was lightly sheared at ∼75 Kbp target length for library construction using PacBio SMRTbell Express Template Prep Kit 2.0. The resulting library was selected for inserts approximately greater than 25 Kbp with the BluePippin (Sage Science) and sequenced on one SMRT cell 8M on Sequel II for 30 hours of data capture.

### Hi-C library and sequencing

A Hi-C library was constructed from liver DNA of the same *T. borchgrevinki* individual by Phase Genomics, Inc. using their Proximo Hi-C kit. The restriction nuclease DpnII was used for chromatin fragmentation. The Hi-C library was quantified by qPCR and then sequenced by Phase Genomics on an Illumina NovaSeq6000 machine to generate 2×150bp paired-end reads.

### Generation of *de novo* contig- and chromosome-level genome assemblies

To generate a *de novo* contig-level genome assembly of *T. borchgrevinki*, we processed our raw CLR data using two different strategies and then assembled the respective data sets with two different assemblers (each of which applied a complementary assembly algorithm).

In our first raw data preparation, we processed the CLR long-reads to remove potential chimeras. We aligned raw reads to each other using `minimap2` (v2.1; Li (2018)) with an all-versus-all approach (using PacBio preset `ava-pb` and mapping option `-g 5000` to set a maximum distance between seeds for overlap generation). We then used `fpa` (v0.5.1; Marijon et al. (2020)) to filter alignments if the overlaps had a length less than 2,000 bp (`-- length-lower 2000`) or they were formed by an internal match, i.e., all the nucleotides in one read were contained in a second read (`--internalmatch`). Next, we used `yacrd` (v0.6.2; Marijon et al. (2020)) to remove reads detected as chimeric and reads that contained significant regions with low coverage (coverage ≤3x across 40% or more of the length of the read; `--coverage 3 --not-coverage 0.4`) to create the final raw read data set.

In our second raw data preparation approach, we used a custom Python script, `sample_reads.py` (Rayamajhi *et al*. 2022) to select a subset of reads bounded by a read length minimum, maximum, and total basepair coverage. With this script, we sampled raw CLR reads between 10 and 40 Kpb in length, for a total coverage of 70 Gigabasepairs (Gbp) given an approximately 1Gbp genome size.

These two complementary read sets were separately assembled using `Flye` (v2.6; Kolmogorov et al. (2019)) and `WTDBG2` (v2.5; Ruan and Li (2020)). We further polished the `WTDBG2` assemblies with the raw CLR reads using the `arrow` module in `GCpp` (v2.0.0; Pacific Biosciences).

We estimated contiguity statistics for the `Flye` and `WTDBG2`-based assemblies using `QUAST` (v4.6.2; Gurevich *et al*. (2013)) and based on those metrics, we retained the `Flye` assembly constructed from the first, chimera-detection strategy, and we retained the `WTDBG2` assembly based on the second, subsampling strategy. However, we considered the `Flye` assembly to be our primary assembly, as it displayed an optimal set of assembly metrics (scaffold N50, contig number, total length). Both assemblies were retained into the next stage of analysis so that assembly results could be compared between them to make manual, synteny-based corrections.

To further scaffold the two assemblies, we aligned Hi-C reads to them and generated lists of Hi-C contacts using `Juicer` (v1.6.2; Durand *et al*. (2016)). Each list of Hi-C contacts and its corresponding contig-level assembly were fed to `Juicer`’s `3d-DNA` pipeline for ordering, orienting, and joining the contigs to produce chromosome-level scaffolds. Moreover, for each scaffolded assembly, the information on structural constituents of chromosomes (i.e., description of contigs or scaffolds organized in chromosomes) was exported into an AGP file using a custom Python script.

We assayed the chromosome-level assemblies with `QUAST` and we then assessed them for completeness using `BUSCO` (v5.1.3; Simão *et al*. (2015)) along with the 3,640 Actinopterygii-specific single-copy ortholog genes (`actinopterygii_odb10`) using default parameters.

### Repeat annotation techniques

For each assembly, we identified and masked repetitive sequences. We first generated a *de novo* custom repeat library using `RepeatModeler` (v2.0.4; Flynn *et al*. (2020)). We combined this *de novo* library with the known repeat library for teleost fishes, which we obtained from `Repbase` (Bao *et al*. 2015). This pooled library was then used to identify and soft mask repetitive elements using `RepeatMasker` (v4.1.5; Smit, Hubley & Green, *RepeatMasker Open-4*.*0*). From the annotated repeats, we then calculated the repeat landscape of the genome using the `RepeatMasker calcDivergenceFromAlign.pl` script. Finally, we used a custom Python script to calculate the density of repeat elements along the genome, by tallying the number of annotated repeat elements within a sliding window (calculating a new 250 Kbp window every 100 Kbp).

### Gene finding methods

For gene annotation, RNAseq reads were retrieved from a previously published study of *T. borchgrevinki* (SRA accession SRP018876; Bilyk and Cheng (2014)). These RNAseq reads were mapped to each masked assembly using `STAR` (v2.7.1.a; Dobin *et al*. 2013). The RNAseq alignments were combined with the set of aligned zebrafish proteins (obtained from `OrthoDB` (v11; Kuznetsov *et al*. (2023)) and the `BRAKER3` (Brůna *et al*. 2021) pipeline was executed to annotate genes. The gene predictions from `BRAKER3` were processed using `TSEBRA` (Gabriel *et al*. 2021) to retain gene annotations best supported by either proteins and/or transcripts. The curated genes were annotated for their functions using `InterProscan` (Quevillon *et al*. 2005). The names of genes obtained from the functional annotation analysis were retained. Like repetitive elements, we calculated the density of annotated protein-coding genes along the genome, by summing the number of annotated genes withing a 250 Kbp sliding window (updated every 100 Kbp) using a custom Python script.

### Manual curation via conserved synteny analysis

We used `Synolog` (Catchen *et al*. 2009; Small *et al*. 2016) to identify conserved synteny between genomes to analyze the *T. borchgrevinki* assemblies and to make manual corrections by comparing our `Flye`- and `WTDBG2`-based assemblies against one another. We also expanded this comparison to other notothenioid genomes, including the basal non-Antarctic notothenioid outgroup *Eleginops maclovinus* (NCBI accession GCF_036324505.1; (Cheng *et al*. 2023), and the white-blooded mackerel icefish, *Champsocephalus gunnari* (NCBI accession GCA_036324595.1; Rivera-Colón *et al*. 2023). `Synolog` first identifies orthologous genes by finding reciprocal best `BLAST` hits between the annotated protein-coding genes of two assemblies. The location of orthologs in each genome is determined according to the genome annotation coordinates (specified in the form of a GTF/GFF file). `Synolog` then builds and refines clusters of conserved synteny within chromosomes by implementing a sliding window algorithm to locate and expand conserved gene neighborhoods spanning multiple orthologs. These conserved synteny patterns are then used to define orthology between chromosomes/scaffolds.

We used this synteny-based system to curate our primary assembly (generated by `Flye`). For example, we identified cases in which the Hi-C data was unable to integrate a contig within a larger set of scaffolds, or examples of contigs scaffolded in an incorrect orientation. We then used a set of custom Python scripts to make changes in the order and orientation of these contigs to correct the assembly – in this case bringing the assembly into agreement with orthologous regions of the comparator genomes. The Python script altered the sequence in the FASTA file, updated the AGP description of the genome, and updated coordinates of any gene annotations to produce the final assembly. In addition to assembly curation, we used the conserved synteny patterns between *T. borchgrevinki* and *E. maclovinus*, the closest sister species to cryonotothenioids and the best extant ancestral proxy, to identify and name orthologous chromosomes. Finally, the annotation procedure was re-executed after manual curation to arrive at the final set of genes.

### Results and Discussion

The HMW *T. borchgrevinki* DNA yielded 181.42 Gbp of long-read CLR data, consisting of 7,651,558 reads with a mean length of 23.7 Kbp and a raw read N50 of 33.4 Kbp (Table 1). Additionally, Hi-C library sequencing resulted in 209.01 million 2×150 bp reads. The final *de novo* assembly had a total length of 935.13 Mbp, and was composed of 2,095 scaffolds, with a scaffold N50 of 42.80 Mbp. The assembly contained 23 putative chromosomes (Fig. 1), ranging in size from 17.82 to 65.34 Mbp. The total length of these putative chromosome-level scaffolds was 912.2 Mbp, encompassing 97.55% of the total assembly length. Moreover, 3,523 out of 3,640 BUSCO orthologs were complete (96.8%) and of those complete, 3,481 (95.6%) were found in a single copy, while only 42 (1.2%) were duplicated. Genome assemblies may sometimes exhibit a high count of complete `BUSCO` genes due to the inadvertent increase in complete but duplicated `BUSCO` genes (Rayamajhi *et al*. 2022). However, in this case, the proportion of `BUSCO` genes with a complete status was notably high and the proportion with a duplicated status was minimal.

**Table 1.** Summary of raw PacBio CLR data for *T. borchgrevinki*.

|  |  |
| --- | --- |
| **Read count** | 7,651,558 |
| **Total Length (Mbp)** | 181,428.53 |
| **Mean Length (bp)** | 23,711.32 |
| **N50 Length (bp)** | 33,463 |
| **Reads >20 Kbp** | 4,132,522 |
| **Reads >50 Kbp** | 492,063 |

**Figure 1.**
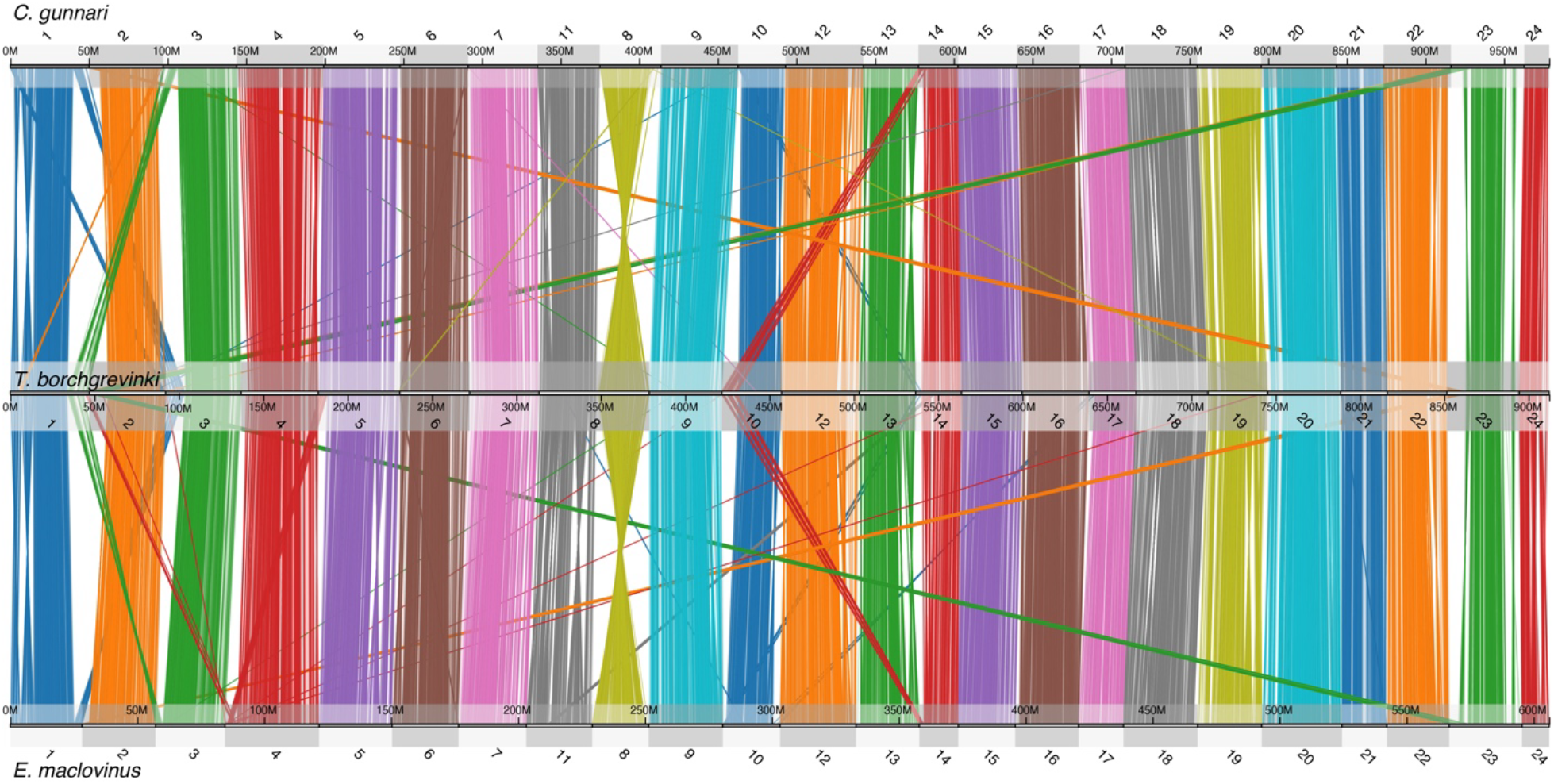
Genome-wide conserved synteny between three notothenioid fishes; the white-blooded mackerel icefish (C. gunnari), the cryopelagic bald notothen (T. borchgrevinki), and the basal, non-Antarctic Patagonian Blennie (E. maclovinus). Each line represents an orthologous gene between two genomes. Clusters of orthologous genes are colored according to their chromosome of origin in each query species, C. gunnari and E. maclovinus for the top and bottom comparisons, respectively.

The total number of predicted protein-coding genes was 22,567 (Table 2), which is comparable to those in the assemblies of other notothenioids (ranging between 20-29 thousand genes) (Bargelloni *et al*. 2019; Bista *et al*. 2020, 2023; Rivera-Colón *et al*. 2023; Cheng *et al*. 2023) suggesting that our annotation process effectively captured the *T. borchgrevinki* protein-coding genes.

**Table 2.** Summary of the final Flye-based genome assembly and BUSCO statistics for *T. borchgrevinki*.

| **Genome assembly statistics** |  |
| --- | --- |
| **Number of chromosome-level scaffolds** | 23 |
| **Total scaffold length** | 935,132,242 |
| **Number of fragments** | 2,095 |
| **Scaffold N50** | 42,797,374 |
| **Scaffold L50** | 10 |
| **Largest Scaffold** | 65,335,880 |
| **Total bases in chromosomes** | 912,238,985 |
| **Percentage of Assembly in chromosomes** | 97.55% |
| **Protein-coding genes** | 22,567 |
| **BUSCO statistics** |  |
| **Complete** | 3,523 (96.8%) |
| **Complete and single copy** | 3,481 (95.6%) |
| **Complete and duplicated** | 42 (1.2%) |
| **Fragmented** | 8 (0.2%) |
| **Missing** | 109 (3.0%) |
| **Total** | 3,640 |

Repetitive elements made up a large proportion of the *T. borchgrevinki* genome. Table 3 shows the breakdown of the interspersed repeats due to DNA transposons, retroelements, SINEs, LINEs, LTRs, and unclassified elements both in terms of absolute length and proportion of the genomes. The total repeat content of *T. borchgrevinki* (54.75%) was 1.6x higher than that of *E. maclovinus* (33.43%; Cheng *et al*. (2023)), and comparable to that of *C. gunnari* (59.45%; Rivera-Colón *et al*. (2023)), another cryonotothenioid species. Repeat classification counts were dominated by DNA transposons, followed by retroelements, and in both cases the length and percentage of these types of repeats followed the same pattern as the total repeat contents among the three species of fish. On a broader scale, the total repeat content of 16 notothenioid species (including three non-Antarctic and 13 cryonotothenioids) ranged from 13% to 54% (Bista *et al*. 2023), which is consistent with our results and places *T. borchgrevinki* on the higher side of repeat content. Moreover, the *T. borchgrevinki* genome displayed a comparatively recent expansion in repeat elements (Fig. 2), a feature conserved with other cryonotothenioids and thought to be strongly associated with the cryonotothenioid radiation and the expansion in genome sizes across the clade (Detrich *et al*. 2010; Auvinet *et al*. 2018; Bista *et al*. 2023).

**Table 3.** Genome length and percentage of interspersed repeats of three notothenioids.

| **Repeat Type** | ***E. maclovinus*** | ***T. borchgrevinki*** | ***C. gunnari*** |
| --- | --- | --- | --- |
| **Assembly length** | 606,289,673 | 935,132,242 | 994,201,109 |
| **DNA transposons** | 96,041,897 (15.84%) | 217,956,285 (23.31%) | 259,784,750 (26.13%) |
| **Retroelements** | 52,256,784 (8.62%) | 146,197,723 (15.63%) | 203,333,903 (20.45%) |
| **SINE** | 389,009 (0.64%) | 7,438,542 (0.80%) | 5,865,787 (0.59%) |
| **LINEs** | 30,658,361 (5.06%) | 78,131,286 (8.36%) | 113,140,086 (11.38%) |
| **LTR elements** | 17,708,326 (2.92%) | 60,627,895 (6.48%) | 84,407,674 (8.49%) |
| **Unclassified** | 34,694,488 (5.72%) | 105,043,501 (11.23%) | 86,396,076 (8.69%) |

**Figure 2.**
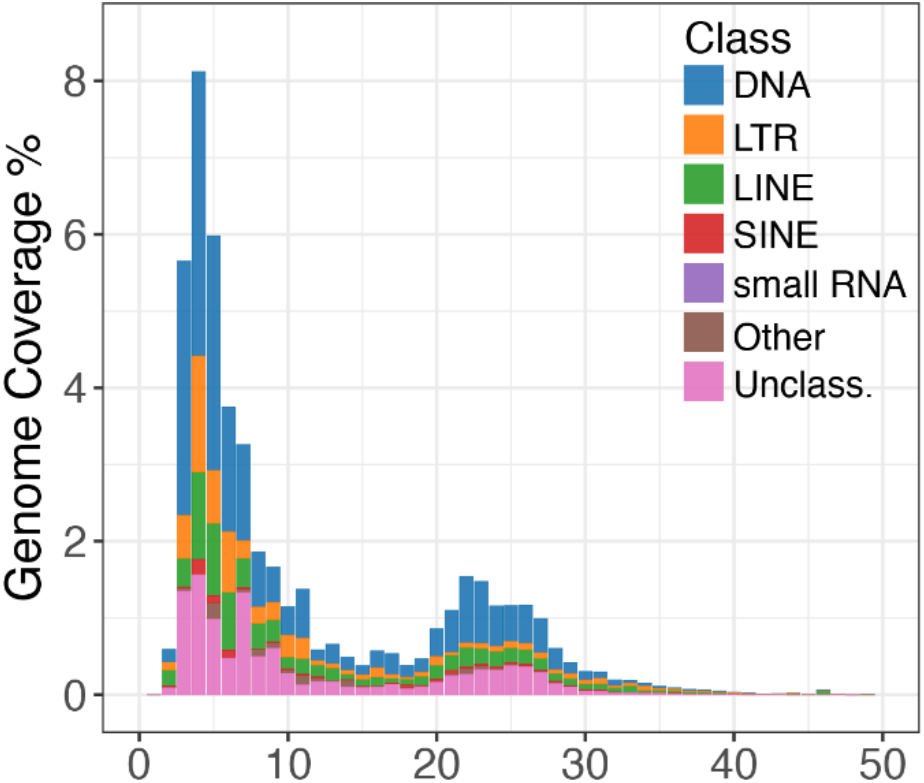
Landscape of repeat divergence in the T. borchgrevinki genome. The X-axis describes Kimura divergence when compared to a consensus element (a proxy for time), while the Y-axis describes coverage of the genome. Colors denote different repeat elements. The T. borchgrevinki genome displays an elevated level of low-divergence repeats, reflecting a relatively recent repeat element expansion.

Genes and repetitive elements appear to have a biased and inversely related distribution in the *T. borchgrevinki* genome (Fig. 3). While the density of protein-coding genes is highest in chromosome centers (Fig. 3A), interspersed repetitive elements appear at a higher density at the terminal ends of chromosomes (Fig. 3B). This distribution of repeats along the chromosomes is similar to two icefishes, *C. gunnari* and *C. esox* (Rivera-Colón *et al*. 2023), and suggests that it might be conserved across members of the Antarctic clade.

**Figure 3.**
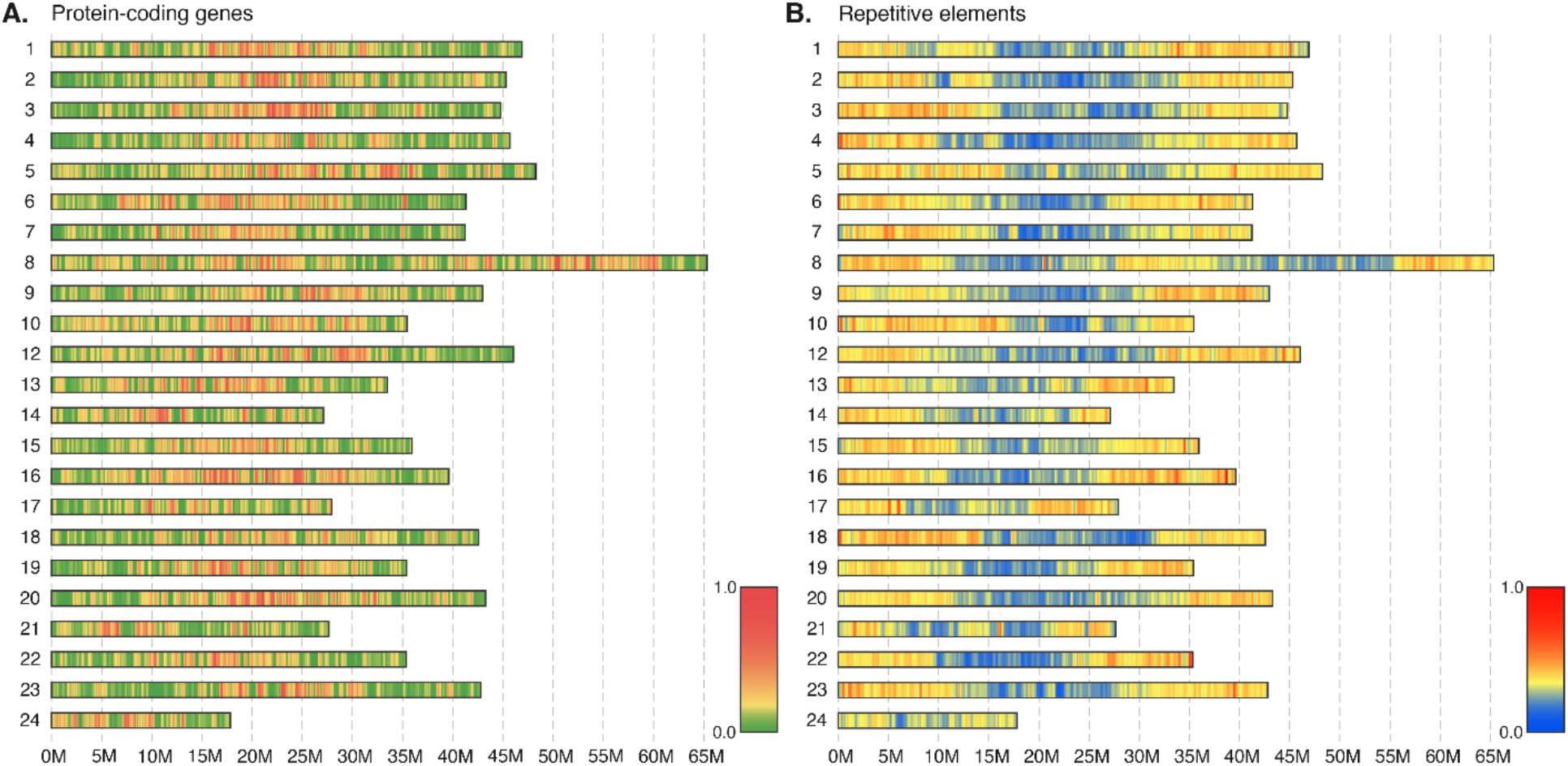
Distribution of genomic elements in the T. borchgrevinki assembly calculated in 250Kbp windows. (A) Protein-coding genes show a higher relative density in chromosome centers. (B) Repeat elements show a higher relative density at the terminal ends of chromosomes. Of particular interest is chromosome 8, formed from the fusion of two ancestral notothenioid chromosomes. Note that element densities are separately calculated for A and B and are thus not directly comparable.

### Conserved synteny analysis reveals *T. borchgrevinki*-specific chromosomal evolution

When we compare the large-scale genomic architecture of the *T. borchgrevinki* genome with the basal, non-Antarctic notothenioid *E. maclovinus* and with the highly derived, white-blooded, Antarctic notothenioid *C. gunnari*, conserved synteny reveals a largely preserved structure consisting of 23 chromosomes (Fig. 1). Our findings are in agreement with previous publications which highlight the large-scale conservation in the karyotype (n = 24) between basal temperate and Antarctic notothenioids (Kim *et al*. 2019; Rivera-Colón *et al*. 2023; Bista *et al*. 2023). However, the *T. borchgrevinki* genome has 23 chromosomes instead of the typical 24, as chromosomes 8 and 11 have fused in *T. borchgrevinki*, resulting in a 65.34 Mbp chromosome which we have continued to label as chromosome 8 (Figs. 1 and 4). This fusion was first shown via karyological analysis in Morescalchi *et al*. (1992) and later in Auvinet *et al*. (2020). With 23 chromosomes, our assembly of a female individual is consistent with the karyological data, as the diploid chromosome number of *T. borchgrevinki* depends on sex (i.e., 2n=45 for males and 2n=46 for females) (Morescalchi *et al*. 1992; Auvinet *et al*. 2020). This centromeric fusion is likely relatively recent, since the structure of the two fused chromosomes remains largely intact, as reflected in the gene and repeat architectures – each sub chromosome shows high gene density in the center and high repeat density on the ends (Fig. 3), while the sub chromosomes show strongly conserved synteny to other notothenioid genomes (Fig. 4).

**Figure 4.**
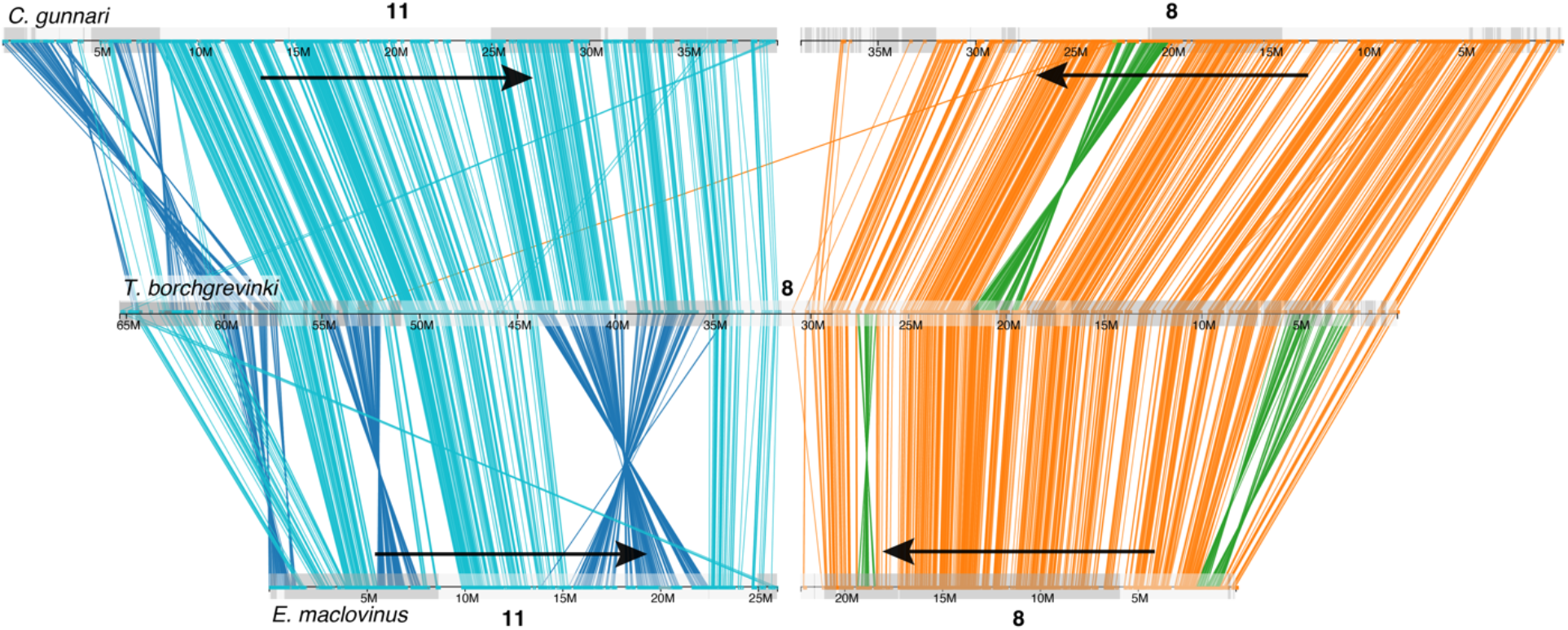
Evidence of a chromosomal fusion in T. borchgrevinki. Conserved synteny plot showing orthology between chromosomes 8 and 11 in C. gunnari (top), the fused chromosome 8 in T. borchgrevinki, and chromosomes 8 and 11 in E. maclovinus (bottom). Colored lines represent orthologous genes between the chromosomes, colored in blue and orange for their ancestral color of origin (11 and 8, respectively). Dark blue and green lines show lineage-specific inversions between the species.

More broadly, at least two notothenioid genera, *Notothenia* and *Trematomus* have major chromosomal fusions. While *Notothenia*’s rearrangements appear to be conserved across the genus, changes among *Trematomus* appear to be independently derived and often species-specific (Amores *et al*. 2017; Auvinet *et al*. 2020). *Trematomus* additionally contains several species (e.g., *T. hansoni* and *T. newnesi*) with sex-specific chromosome numbers identical to *T. borchgrevinki – where females possess a diploid X*_*1*_*X*_*1*_*X*_*2*_*X*_*2*_*configuration but males possess X*_*1*_*X*_*2*_*Y (leading to 2n=46/45, female/male)* – as well as widely different, sex-specific chromosomes numbers (*T. nicolai*; 2n=58/57, female/male) (Morescalchi *et al*. 1992), and even species with intraspecific variable numbers of chromosomes (*T. Ioennbergii;* 2n=26-33) (Morescalchi *et al*. 1992; Ghigliotti *et al*. 2015).

Beyond the fused chromosome 8, there are several large-scale structural variants including translocations and inversions (Fig. 5). Specifically, on six *T. borchgrevinki* chromosomes (1, 2, 3, 10, 14, and 23) we see translocations on the order of 10 Mbp. Interestingly, these inversions appear as swaps of chromosome ends (e.g., chromosomes 1 and 3 have swapped telomeres, as have chromosomes 2 and 23, etc.). In addition, there are several multi-megabase inversions that are specific to this clade (e.g., chromosomes 1, 13 and the fused chromosome 8; Figs. 5 and 4).

**Figure 5.**
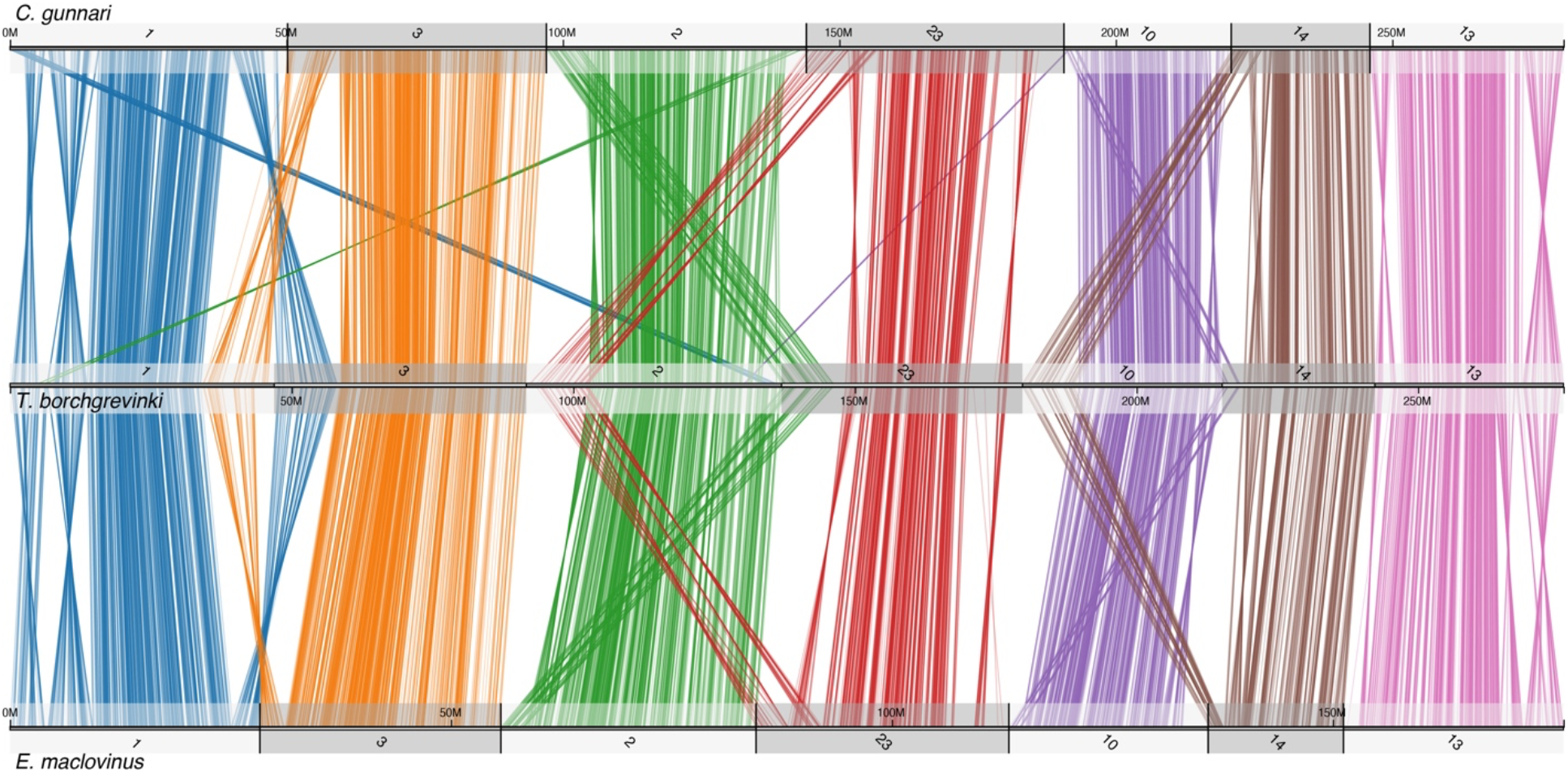
Conserved synteny of seven orthologous chromosomes between three notothenioid fishes highlighting major translocations and inversions that have occurred on the T. borchgrevinki lineage. Top: C. gunnari; middle: T. borchgrevinki; bottom: E. maclovinus. Chromosomes have been ordered (1, 3, 2, 23, 10, 14, and 13) to highlight translocations.

As a major, cold adapted and cold specialized notothenioid, possessing a robust array of antifreeze proteins, this well-constructed assembly holds great potential for facilitating genome-based research in polar and non-polar notothenioids.

## Data Availability

The raw data alongside the assembly and annotation are available in NCBI under BioProject PRJNA907802. Raw PacBio and Hi-C reads can be also found in the NCBI Sequence Read Archive under accessions SRX18476836 and SRX24931301, respectively. Genome assembly and annotation files are also available in Dryad under DOI 10.5061/dryad.h44j0zpv7.

## Conflict of Interest

The authors declare no conflicts of interest.

## Author Contributions

C-HCC collected the specimen and led the molecular work. NR, JC, AGR-C, and BFM performed the analysis. NR, JC, AGR-C, and C-HCC wrote the manuscript. All authors approved the final manuscript.

## Funder Information

Work was supported by the NSF DGE 10-69157 IGERT Grant to AGR-C, by NSF OPP Grant 1645087 to JC and C-HCC, and NSF ANT Grant 11-42158 to C-HCC.

## Notes

### Competing Interest Statement

The authors have declared no competing interest.

### Summary of Updates

The manuscript was updated to correct several typographical errors and to clarify several points in the Introduction and Results.

